# Discovery of Cancer Driver Long Noncoding RNAs across 1112 Tumour Genomes: New Candidates and Distinguishing Features

**DOI:** 10.1101/065805

**Authors:** Andrés Lanzós, Joana Carlevaro-Fita, Loris Mularoni, Ferran Reverter, Emilio Palumbo, Roderic Guigó, Rory Johnson

## Abstract

Long noncoding RNAs (lncRNAs) represent a vast unexplored genetic space that may hold missing drivers of tumourigenesis, but few such “driver lncRNAs” are known. Until now, they have been discovered through changes in expression, leading to problems in distinguishing between causative roles and passenger effects. We here present a different approach for driver lncRNA discovery using mutational patterns in tumour DNA. Our pipeline, ExInAtor, identifies genes with excess load of somatic single nucleotide variants (SNVs) across panels of tumour genomes. Heterogeneity in mutational signatures between cancer types and individuals is accounted for using a simple local trinucleotide background model, which yields high precision and low computational demands. We use ExInAtor to predict drivers from the GENCODE annotation across 1112 entire genomes from 23 cancer types. Using a stratified approach, we identify 15 high-confidence candidates: 9 novel and 6 known cancer-related genes, including *MALAT1*, *NEAT1* and *SAMMSON*. Both known and novel driver lncRNAs are distinguished by elevated gene length, evolutionary conservation and expression. We have presented a first catalogue of mutated lncRNA genes driving cancer, which will grow and improve with the application of ExInAtor to future tumour genome projects.

## Introduction

Whole genome sequencing makes it possible to comprehensively discover the mutations, and the mutated genes, that are responsible for tumour formation. By sequencing pairs of normal and tumour genomes from large patient cohorts, projects such as the ICGC (International Cancer Genome Consortium) and TCGA (The Cancer Genome Atlas) aim to create definitive driver mutation catalogues for all common cancers (1, 2). Focussing on entire genomes, rather than just captured exomes, these studies hope to identify driver elements amongst the ~98% DNA that does not encode protein. These noncoding regions contain a wealth of regulatory sequences and non-coding RNAs whose role in cancer has been neglected until now (3).

Amongst the most numerous, yet poorly understood of the latter are long noncoding RNAs (lncRNAs). These are long RNA transcripts that share many characteristics of mRNAs, with the key difference that they do not contain any recognizable Open Reading Frame (ORF), and thus are unlikely to encode protein (4). LncRNAs perform a diverse range of regulatory activities within both the nucleus and cytoplasm by interacting with protein complexes or other nucleic acids (5). While their expression tends to be lower than protein-coding mRNAs, lncRNAs are thought to be highly expressed in a subset of cells in a population (6). The number of lncRNA genes in the human genome is still uncertain, but probably lies in the range 20,000 - 50,000 (7, 8). This vast population of uncharacterized genes likely includes many with novel roles in cancer.

In recent years a small but growing number of lncRNA have been implicated in cancer progression through various mechanisms (9). LincRNA-P21, a tumour suppressor, acts downstream of P53 by recruiting the repressor hnRNP-K to target genes (10). Proto-oncogene lncRNAs include HOTAIR, upregulated in multiple cancers, which recruits the repressive PRC2 chromatin regulatory complex to hundreds of genes (11). Cancer-related lncRNA have features of functional genes, including sequence conservation, orthologues in other mammals, chromatin marks and regulated subcellular localisation (4). Moreover they display typical characteristics of cancer drivers, including influence on cellular phenotypes of proliferation and apoptosis, and in clinical features such as patient survival and altered expression across tumour collections (3, 8, 11).

The absence of whole-genome maps of somatic mutations has meant that searches for new cancer-related lncRNAs have relied on conventional transcriptomic approaches that reveal changes in their expression levels that accompany cancer. However such approaches are not capable of distinguishing passenger and driver effects, nor do they identify mutations in the mature lncRNA sequence that may drive tumourigenesis independent of upstream regulatory changes (8, 12, 13). Two recent studies clearly demonstrate that somatic mutations, in these cases amplifications of entire loci, can drive tumour formation (14, 15). Nevertheless, we remain largely ignorant of the role that mutations in lncRNA genes play during the early stages of tumourigenesis.

The statistical analysis of somatic mutation patterns is a powerful means of identifying genes that drive early tumour formation. For protein-coding genes, tools have been developed to successfully identify mutations that result in gain or loss of function in translated peptide sequences (16). These tools take advantage of exome sequencing – the targeted capture and sequencing of approximately 2% of the genome encoding protein (17). A number of methods applied to this problem for protein-coding genes (16) take advantage of the fact that cancer-associated genes (both tumour-suppressors and proto-oncogenes) display characteristic and non-random mutational patterns in their protein-coding sequence. They prioritise genes with mutations that are predicted to result from positive selection on the encoded protein.

As such, these methods are generally inapplicable to lncRNA, which do not encode protein, and for which we presently do not have maps of their functional sequence domains, nor an understanding of the molecular mechanism of such domains. While capable of discovering protein-coding driver genes, exome sequencing ignores mutations occurring in the multitude of noncoding regulatory elements known to exist in the human genome (18). Noncoding driver mutations can be comprehensively discovered for the first time by projects to sequence collections of entire cancer genomes (1). In the present study, we describe and characterise a tool, called ExInAtor, for the discovery of driver lncRNA genes. ExInAtor identifies genes with excess of exonic mutations, compared to the expected local neutral rate estimated from intronic and surrounding sequences. We present a comprehensive prediction of candidate lncRNAs across 1104 genomes from 23 cancer types. These candidates have a series of features consistent with their being genuine drivers.

## Results

### A method for discovering driver genes from cancer genomes

Our aim was to develop a method to identify tumour driver long noncoding RNAs (lncRNAs) using short nucleotide variant (SNV) mutations from cancer genome sequencing projects. We define SNVs, from now on, as somatic substitutions or indels of length 1 nt. The majority of lncRNAs are spliced, and we assume throughout that their functional sequence resides in exonic regions that are incorporated into the mature transcript (19). Intronic sequence is removed during splicing and hence is not directly relevant to their function. Consequently, we hypothesised that driver lncRNAs will display an excess of somatic mutations in exons compared to the local background mutational rate, estimated by their introns and flanking genomic regions – henceforth referred to as “background regions”. This approach is conservative, given that background regions are likely to include functional regulatory elements that may themselves carry driver mutations.

We implemented this approach in a computational pipeline called ExInAtor (Fig. 1 and File S1). ExInAtor requires two principal inputs: an annotation of lncRNA genes and a catalogue of tumour mutations. At its heart, ExInAtor employs a parametric statistical test to identify genes that present a significantly elevated exonic mutation rate compared to local background regions. The latter are comprised of intronic and flanking genomic sequence. We took care to account for a key confounding factor: the unique mixture of mutational signatures that characterises every individual tumour, and every tumour type (20). Such signatures can be described as a probability for every nucleotide to mutate to every other, conditioned on the identity of flanking positions - summarised in a matrix of 96 trinucleotide substitution frequencies (20). In other words, mutation rates are dependent on nucleotide composition. The mutational signature must be taken into account when comparing mutational loads of exons to surrounding regions, because they tend to have marked differences in nucleotide composition - both for protein-coding genes and lncRNAs (21).

**Fig 1:**
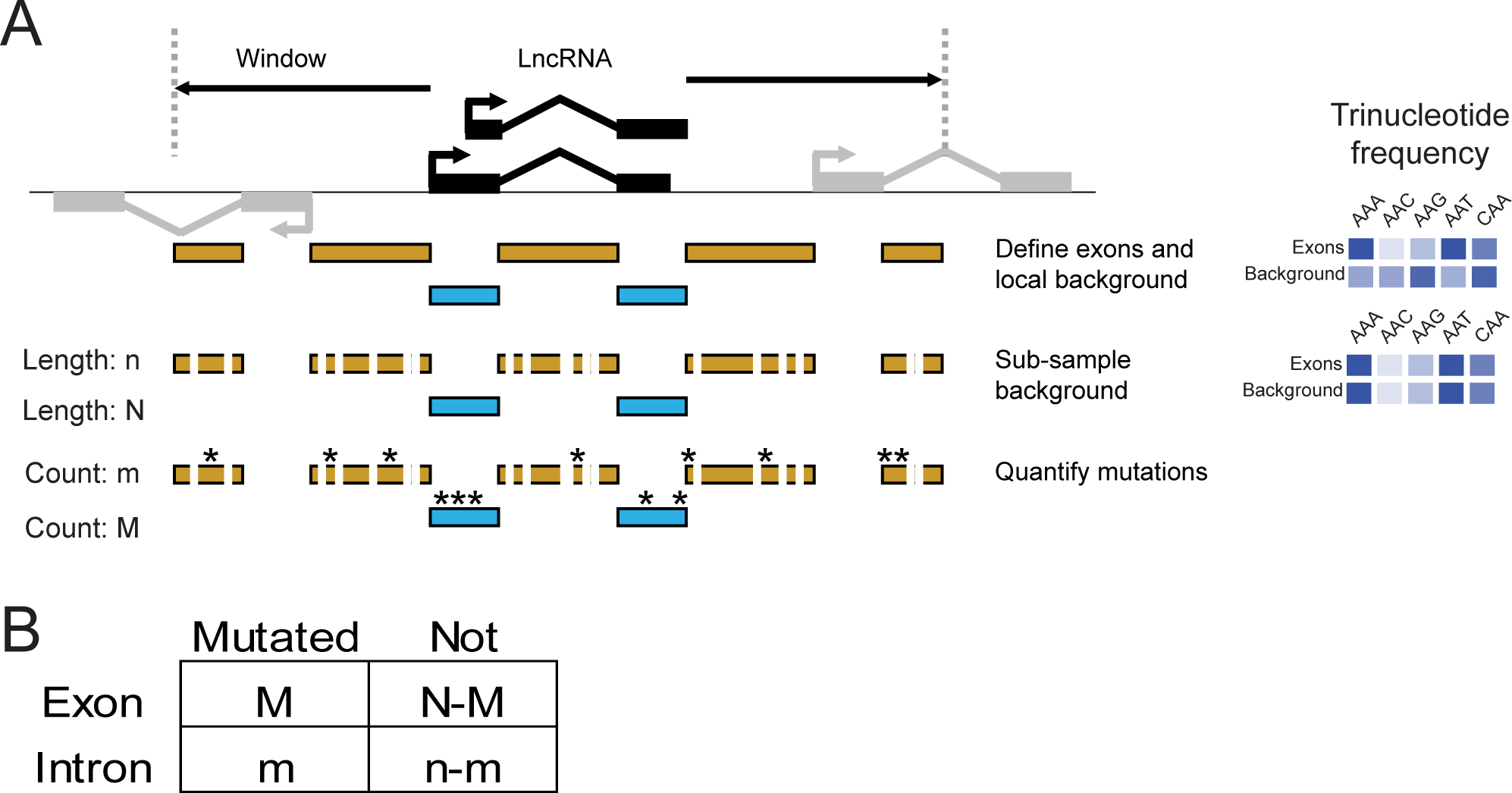
Outline of the ExInAtor method. (A) The steps of gene definition, subsampling and analysis performed to quantify exonic and background mutations. Sampling is performed in such a way that, at the end, the trinucleotide frequency of the background region is identical to the exonic region. (B) The number of mutations in background and exonic regions is compared by a contingency table analysis.

ExInAtor employs a subsampling approach to balance the trinucleotide content of exons and reference regions, thereby accounting for mutational signatures (Fig. 1A). Exonic regions of each gene are defined as the projection of all exons from the union of its transcripts. Next, the reference region is defined as all non-exonic nucleotides within the gene, in addition to upstream and downstream windows of defined length. Within these exonic and reference regions, the frequencies of trinucleotides are calculated. Then, nucleotides are randomly sampled (without replacement) from the reference region, until the maximum possible amount of sequence with identical trinucleotide composition has been collected. Now, the number of SNVs overlapping exons, M, and those overlapping remaining reference nucleotides, m, are compared using a contingency-table analysis and statistical significance is calculated according to hypergeometric distribution (Fig. 1B) (see Materials and Methods for more details).

We prepared a carefully-filtered lncRNA annotation, to avoid several potential sources of false positive predictions. We were particularly concerned by two potential confounding factors: first, misinterpretation of mutations that may affect protein-coding regions overlapping the same DNA as lncRNA exons; and second, the presence of mis-classified protein-coding transcripts among the GENCODE annotation (4). Thus, we removed genes of uncertain protein-coding potential, as judged by computational protein-coding potential classifiers (see Materials and Methods). We also removed any lncRNA genes, such as cis-antisense and intronic lncRNAs, that overlap annotated protein-coding genes. In this way we narrowed the set of GENCODE v19 lncRNA genes from 13,870 to 5,887 intergenic, confidently-noncoding lncRNAs (Table 1). To this set we added back 27 cancer-related, GENCODE v19 lncRNAs from the literature (see below).

**Table 1:** Filtered gene sets.

| **Element** | **LncRNA** |  |  | **Protein coding** |  |  |
| --- | --- | --- | --- | --- | --- | --- |
|  | **CRL** | **Non-CRL** | **Total** | **CGC** | **Not CGC** | **Total** |
| Genes | 45 | 5,869 | 5,914 | 545 | 19,769 | 20,314 |
| Transcripts | 297 | 9,086 | 9,383 | 3,239 | 78,463 | 81,702 |
| Exons | 1,259 | 27,025 | 28,284 | 35,902 | 702,974 | 738,876 |
| Merged Exons | 267 | 19,153 | 19,420 | 9,326 | 218,186 | 227,512 |

One advantage of ExInAtor is its indifference to genes’ biotype. This arises from its lack of reliance on measures of functional impact(22), meaning that it can equally be used on lncRNAs or protein-coding genes. Indeed, similar approaches have been used to discover coding driver genes in the past (23). We took advantage of this to assess its ability to discover known protein-coding driver genes from the Cancer Gene Census (24) amongst the GENCODE annotation. This provided us with a useful independent validation of ExInAtor’s precision, of particular value given the low number of known driver lncRNAs at present.

### Datasets of somatic mutations in cancer genomes

To search for lncRNA driver genes, we took advantage of the two largest available sources of cancer genome mutations: one collected by the Cancer Genome Project at the Sanger Institute, hereafter named “Alexandrov” (20), and the other from The Cancer Genome Atlas (TCGA) (1) (Table 2). These data were aggressively filtered to remove potential artefacts arising from germline mutations (see Materials and Methods). The Alexandrov dataset comprises 9 cancers with between 15 and 119 individuals and 10,436 and 2,796,863 mutations each. The TCGA dataset consists of 14 cancers with between 15 and 96 individuals and 21,113 to 4,680,653 mutations each. Of note is the large spread in sample sizes and mutation rates across tumour types. Taking all cancers together, we observed an excess of mutations in lncRNAs over protein-coding genes, and in background over exons, suggesting a general selective pressure against disruptive mutations in both gene classes (File S2).

**Table 2:** Cancer datasets used in this study.

| **Dataset** | **Cancer** | **Mutations** | **Genomes** |
| --- | --- | --- | --- |
| Alexandrov | Breast | 655,823 | 119 |
| Alexandrov | CLL | 51,377 | 28 |
| Alexandrov | Liver | 867,080 | 88 |
| Alexandrov | Lung_adeno | 1,520,078 | 24 |
| Alexandrov | Lymphoma_B-cell | 126,581 | 24 |
| Alexandrov | Medulloblastoma | 123,642 | 100 |
| Alexandrov | Pancreas | 110,944 | 15 |
| Alexandrov | Pilocytic_astrocytoma | 10,436 | 101 |
| Alexandrov | Stad | 2,796,863 | 100 |
| Alexandrov | Pancancer | 6,259,996 | 607 |
| TCGA | BLCA | 385,128 | 21 |
| TCGA | BRCA | 620,238 | 96 |
| TCGA | CRC | 4,680,653 | 42 |
| TCGA | GBM | 180,896 | 27 |
| TCGA | HNSC | 295,709 | 27 |
| TCGA | KICH | 24,508 | 15 |
| TCGA | KIRC | 131,828 | 29 |
| TCGA | LGG | 35,474 | 18 |
| TCGA | LUAD | 1,237,722 | 46 |
| TCGA | LUSC | 1,626,973 | 45 |
| TCGA | PRAD | 21,113 | 20 |
| TCGA | SKCM | 3,538,750 | 38 |
| TCGA | THCA | 37,882 | 34 |
| TCGA | UCEC | 2,268,210 | 47 |
| TCGA | Pancancer | 14,841,279 | 505 |
| Both | Superpancancer | 20,837,263 | 1112 |

### The landscape of driver lncRNAs across 23 tumour types

To comprehensively discover candidate lncRNA drivers, ExInAtor was run on the 23 tumour types described above. We adopted some analysis strategies to account for the relatively shallow nature of the data and our consequently weak statistical power to find driver genes. First, in order to discover both cancer-specific and ubiquitous driver genes, ExInAtor was run on each dataset in distinct configurations: (1) grouping samples by tumour type (“Tumour Specific”), (2) pooling together the entire set of tumours within each of the two projects (“Pancancer”) and (3) pooling data across both projects (“Superpancancer”).

Second, we used sample stratification to boost sensitivity. This approach is commonly used when statistical power is reduced by multiple hypothesis testing (25,26). LncRNA genes were divided into two groups of different sizes, and each was treated independently during multiple hypothesis correction. This reduces the burden on resulting false discovery rate estimates. As a reference set, we curated 45 experimentally-validated cancer-related lncRNAs from the scientific literature, henceforth “Cancer-Related LncRNAs” (CRLs) (File S3). All CRL genes belong to GENCODE v19 annotation. Remaining filtered lncRNAs are referred to as “Non-CRL” (File S4). Summary statistics of the gene sets used are shown in Table 1.

At a Q value (false discovery rate) cutoff of 0.1, we discovered a total of 15 lncRNAs (6 and 9 from CRL and non-CRL, respectively) (Fig. 2A) (Files S5 and S6) and 24 protein-coding genes (File S7). Relaxing the cutoff to Q<0.2, we discover 10 and 27 CRL and non-CRL lncRNAs, respectively. Henceforth we refer to these as driver genes, and a Q-value threshold of 0.1 is assumed unless stated otherwise. ExInAtor predicted a total of five lncRNA driver genes in Alexandrov tumours, nine in TCGA and two in Superpancancer (one of them already detected in Pancancer TCGA). The greatest numbers of drivers predicted in individual tumours were three apiece in Breast and Kidney Chromophore (Fig. 2D).

**Fig 2:**
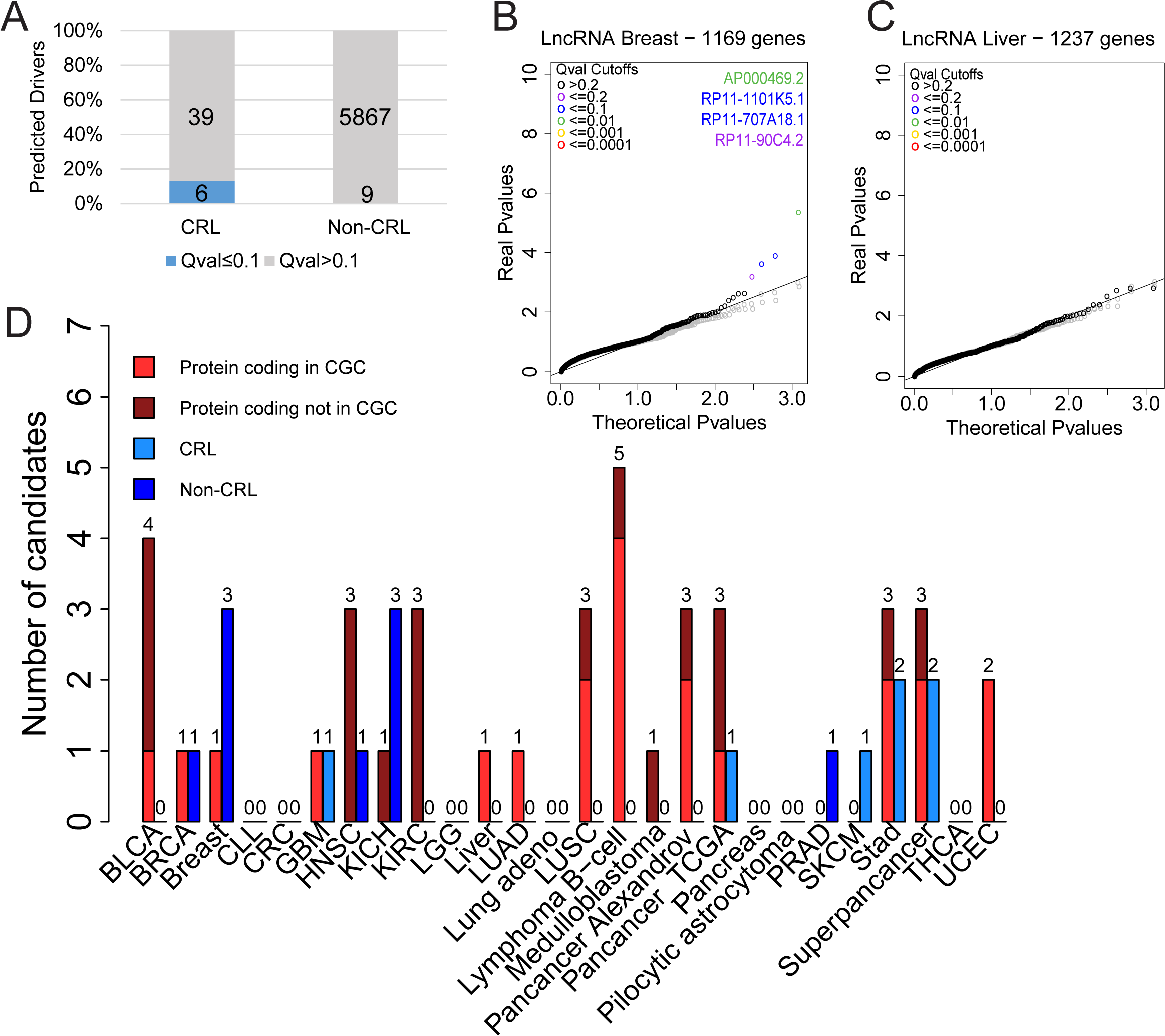
The landscape of driver lncRNAs across 23 tumour types. (A) The numbers and proportion of literature-reported cancer-related long noncoding RNAs (CRL) and non-CRL candidates identified in this study. Q value is equivalent to false discovery rate (FDR). (B) A Quantile-Quantile (QQ) plot showing the performance of ExInAtor on Breast cancer mutations. Each point represents one gene. Note the deviation of real P values from the theoretical distribution in a small tail of cases. Simulated data was created by randomising mutations while maintaining trinucleotide context. (C) As for B, for Liver cancer. Note the lack of candidates in this dataset. (D) The number of driver genes discovered at a Q cutoff of 0.1 across the Alexandrov and TCGA collections. Cancer Gene Census (CGC) are true positive, known protein-coding cancer driver genes.

Several findings suggest that false positive prediction rates are low. Reported P values closely follow the expected null distribution for the majority of genes (a full set of Quantile-quantile (QQ) plots can be found in File S8). Furthermore, while a number of tumour types display a small number of putative driver lncRNAs that strongly deviate from the null expectation (exemplified by Breast cancer sample in Fig. 2B), other samples yield no candidates at all (eg Liver cancer, Fig. 2C). In general, inspection of QQ plots shows a tendency for deflation of P values (File S8). To further test false discovery rates, we reran these analyses on tumour data that had been randomised using two different methods (see Materials and Methods for details). ExInAtor predicted no lncRNA drivers in either dataset (grey dots in Figs. 2B&C and File S8). Together these data point to a rather conservative statistical model, which may discard some *bona fide* drivers. A comprehensive set of predictions across all analyses can be found in File S9.

### ExInAtor identifies known and novel lncRNA driver genes

ExInAtor’s sensitivity is demonstrated by its identification of altogether six CRL genes. These are: *MALAT1, NEAT1, PCA3, BCAR4,* lncRNA-ATB (CTD–2314B22.3) and the recently-discovered melanoma driver *SAMMSON* (RP11–460N16.1) (Table 3). The latter was detected in stomach adenocarcinoma, and we found that it is also present in stomach RNAseq (File S10). The majority of candidates were found in tumour-specific analysis (Fig. 3A). Nevertheless, two CRL lncRNAs, *NEAT1* and *MALAT1,* were identified in Pancancer analysis, consistent with a general role in tumourigenesis: both are long, unspliced and nuclear-retained lncRNAs with demonstrated roles across a range of cancer types (9). As shown in Fig. 3B, the *NEAT1* exon region experiences an elevated mutation rate across cancers, when compared to its flanking background regions. *NEAT1* was identified in a recent study of liver cancer genomes, and as the authors pointed out, it cannot be ruled out that it is identified through increased local mutation rate (27).

**Fig 3:**
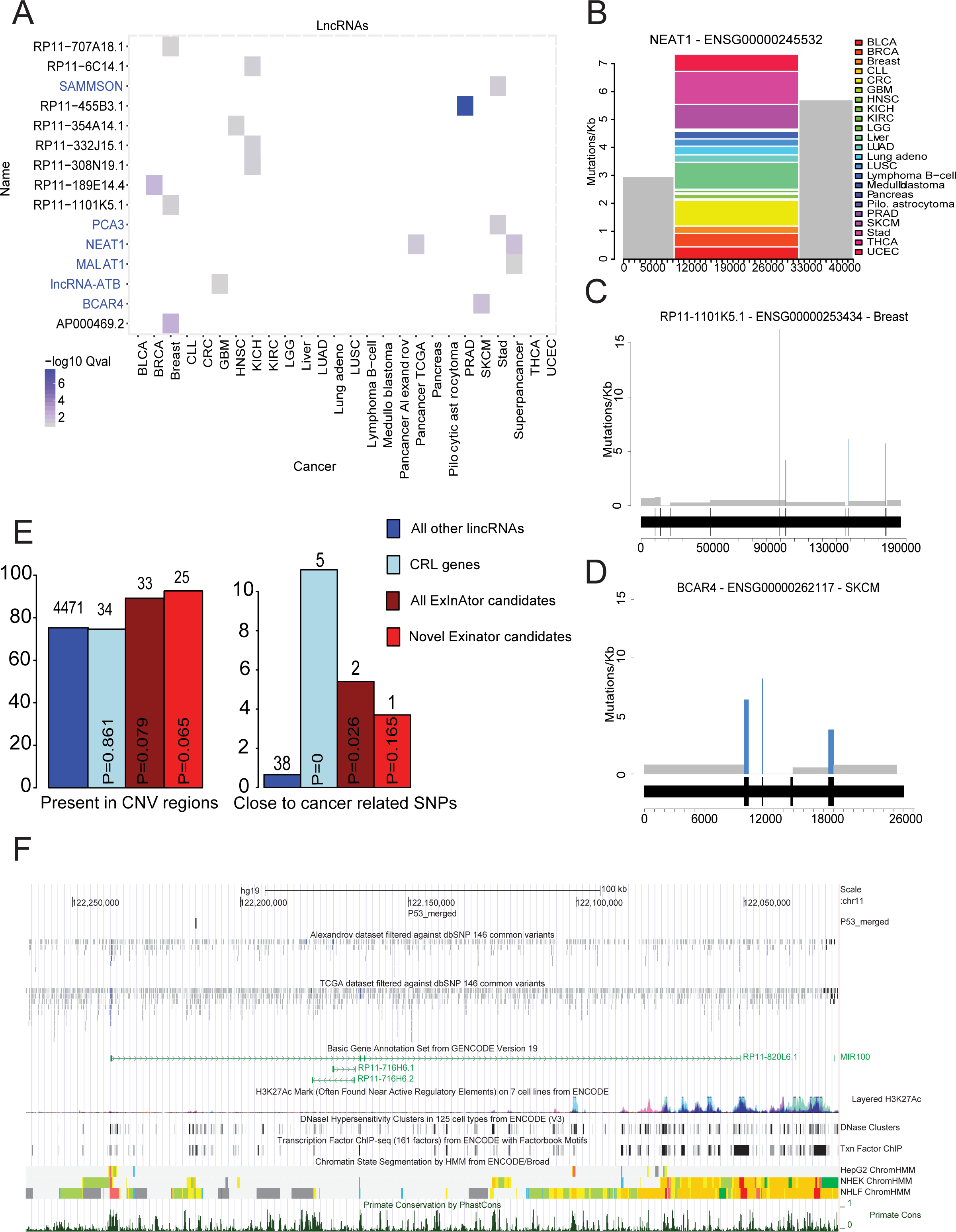
LncRNA cancer driver genes predicted by ExInAtor across cancer genomes. (A) All driver lncRNAs (Q≤0.1) and the tumour type in which they are identified. Gene names in blue indicate those belonging to CRL. (B) A mutation density plot for NEAT1 in all cancers, plotting the SNVs per kilobase as a function of gene regions. Grey represent background regions, while colours represent the mutational contribution of each cancer type to the single exon. The x-axis represents position, in bp, with respect to the start of the background region, defined here to be at 10 kb upstream of the gene’s annotated TSS. (C) The Breast mutation profile of RP11-1101K5.1, a gene with mutations in four exons. Rectangles depict mutational density of exons (blue) and introns (grey). The gene structure is indicated below, where wider portions represent exons, separated by narrower introns. (D) The Breast mutation frequency in *BCAR4.* (E) Percentage of genes and candidates in CNV regions and proximal to cancer-related germline SNPs. Numbers above bars indicate the absolute numbers of genes represented by each percentage. Statistical significance in each case was estimated using Fisher’s Exact test. (F) An example of an ExlnAtor-predicted novel candidate gene, *RP11-820L6.1.* Note the presence of promoter-like histone marks (red, ChromHMM track), evolutionary conservation (PhastCons Primate conservation), and cancer SNVs around the gene TSS, as well as a proximal P53 binding site (“P53_merged”).

**Table 3:** Table 3: List of predicted lncRNA drivers at Q<0.1. Known cancer genes from CRL are marked in bold face. “Pc” - PanCancer.

| Cancer | Gene Name | Gene ID | Ex mut | Ex len | Intr mut | Intr len | Pval | Qval | Ex mut rate | Intr mut rate | Ratio |
| --- | --- | --- | --- | --- | --- | --- | --- | --- | --- | --- | --- |
| Breast | AP000469.2 | ENSG00000224832 | 9 | 238824 | 2 | 1176908 | 4.38E-06 | 5.05E-03 | 3.77E-02 | 1.70E-03 | 22.2 |
| Stad | **PCA3** | ENSG00000225937 | 10 | 392190 | 20 | 2702680 | 2.86E-03 | 5.49E-02 | 2.55E-02 | 7.40E-03 | 3.4 |
| KICH | RP11-308N19.1 | ENSG00000234323 | 1 | 26384 | 1 | 2437289 | 2.13E-02 | 6.39E-02 | 3.79E-02 | 4.10E-04 | 92.4 |
| Stad | **SAMMSON** | ENSG00000240405 | 5 | 208795 | 1 | 697899 | 3.14E-03 | 5.49E-02 | 2.39E-02 | 1.43E-03 | 16.7 |
| GBM | **lncRNA-ATB** | ENSG00000244306 | 4 | 314168 | 4 | 1347296 | 4.68E-02 | 9.36E-02 | 1.27E-02 | 2.97E-03 | 4.3 |
| Super_Pc | **NEAT1** | ENSG00000245532 | 163 | 25316741 | 16 | 5559984 | 4.98E-04 | 2.19E-02 | 6.44E-03 | 2.88E-03 | 2.2 |
| Pc_TCGA | **NEAT1** | ENSG00000245532 | 96 | 11497239 | 7 | 2524993 | 9.28E-04 | 4.08E-02 | 8.35E-03 | 2.77E-03 | 3.0 |
| PRAD | RP11-455B3.1 | ENSG00000248202 | 7 | 16573 | 1 | 380379 | 1.70E-09 | 1.19E-08 | 4.22E-01 | 2.63E-03 | 160.7 |
| KICH | RP11-332J15.1 | ENSG00000249734 | 3 | 8277 | 15 | 314985 | 1.03E-02 | 6.39E-02 | 3.62E-01 | 4.76E-02 | 7.6 |
| Breast | RP11-707A18.1 | ENSG00000250125 | 6 | 485395 | 9 | 6908893 | 2.38E-04 | 9.16E-02 | 1.24E-02 | 1.30E-03 | 9.5 |
| KICH | RP11-6C14.1 | ENSG00000250488 | 1 | 5369 | 1 | 687929 | 1.54E-02 | 6.39E-02 | 1.86E-01 | 1.45E-03 | 128.1 |
| Super_Pc | **MALAT1** | ENSG00000251562 | 83 | 9683213 | 40 | 7768392 | 4.45E-03 | 9.79E-02 | 8.57E-03 | 5.15E-03 | 1.7 |
| Breast | RP11-1101K5.1 | ENSG00000253434 | 6 | 165285 | 19 | 4878981 | 1.28E-04 | 7.38E-02 | 3.63E-02 | 3.89E-03 | 9.3 |
| HNSC | RP11-354A14.1 | ENSG00000254689 | 3 | 12579 | 4 | 697784 | 1.84E-04 | 8.14E-02 | 2.38E-01 | 5.73E-03 | 41.6 |
| BRCA | RP11-189E14.4 | ENSG00000261623 | 9 | 352599 | 1 | 1252991 | 9.53E-06 | 9.25E-03 | 2.55E-02 | 7.98E-04 | 32.0 |
| SKCM | **BCAR4** | ENSG00000262117 | 6 | 50458 | 8 | 533056 | 6.81E-04 | 2.11E-02 | 1.19E-01 | 1.50E-02 | 7.9 |

One important potential source of false positive signal in this study could be elevated mutational rates in DNA regulatory elements, such as enhancers, which happen to overlap the exon of a lncRNA annotation. Such cases would be expected to produce driver lncRNAs, where all mutations are concentrated in a single exon. This would be indistinguishable from *bona fide* driver lncRNAs that have an important functional domain located in a single exon. To investigate this further, we inspected the exon-level mutational density of all candidate lncRNAs (File S11). Intriguingly, we find at least two cases where mutations are elevated across multiple exons, but not intervening introns (Figs. 3C&D). Altogether of 13 multi-exonic candidate lncRNAs, five have mutations in more than one exon. This supports the interpretation that, for these cases at least, mutations cause gain- or loss-of-function in mature lncRNA transcripts, and not through disruption of a DNA-encoded element.

Amongst the novel candidate driver genes were a number of intriguing cases with various characteristics of functionality, none of which have been described in the scientific literature. In Figure 3F we highlight one case, *RP11-820L6.1,* whose promoter is characterised by canonical histone modifications, obvious evolutionary conservation and the recruitment of transcription factors. Most notably, the master tumour suppressor transcription factor and regulator of several cancer lncRNAs, P53, is bound within the first intron (28).

We further sought to establish the degree of overlap between ExInAtor-predicted driver genes and candidates predicted by transcriptomic analyses. Two previous studies to identify cancer-related lncRNAs have searched for differentially-expressed transcripts in cancer transcriptome data from microarrays and RNA sequencing (8,12). From each study we extracted those transcripts that overlap the filtered geneset used here, retrieving a total of 80 and 186 genes from the Du et al and Iyer et al (“MiTranscriptome”) studies, respectively (Files S12 and S13)(8,12). Three genes are identified by both ExInAtor and MiTranscriptome *(PCA3, NEAT1* and *MALAT1)* (P=0.0026, Chi-square with Yates' correction test) and another with Du et al *(PCA3)* (P=0.5, Chi-square with Yates' correction test) (File S14). It should be noted that all these genes belong to the CRL set. MiTranscriptome and Du share 11 genes (P<=0.0001, Chi-square with Yates' correction test). This surprising discordance of driver gene prediction between studies, in addition to their lack of overall intersection with the published CRL set, suggests that (1) these large-scale predictions have considerable false negative rates, and (2) that available catalogues of cancer-related lncRNAs, represented by the CRL set, are incomplete.

We searched for independent evidence of cancer roles for ExInAtor-predicted candidates. Importantly, we separately considered (1) the entire set of candidates, including known CRL genes, and (2), the novel ExInAtor candidates alone. This ensures that findings are not biased by the inclusion of experimentally-verified CRL drivers amongst candidate gene sets. We first tested the frequency with which candidates are affected by copy number variants (CNVs) across matched cancers (29). We found that all candidates, and novel candidates alone, both display a trend to have elevated rates of copy number variation (Fig. 3E). We also investigated whether candidates are more proximal to germline cancer mutations (29). Once more, we observe a trend for candidates to be more likely to be proximally located to such mutations than expected by chance. Although the small numbers involved do not generally reach statistical significance, these findings are additional evidence that ExInAtor predictions, either including or excluding known cancer-related lncRNAs, are involved in tumour progression.

### ExInAtor identifies known protein-coding cancer genes

Although ExInAtor was designed with lncRNAs in mind, it makes no use of functional impact predictions and hence is agnostic to the protein-coding potential of the genes it analyses. We took advantage of this versatility to further test ExInAtor’s precision, by comparing predictions to the gold-standard catalogue of the Cancer Gene Census (CGC) (24). CGC is a manually-curated and regularly-updated annotation of genes whose somatic mutations have been associated with cancer. CGC genes represent a subset of 545 genes (File S15) (2.7%) of the entire GENCODE set of 20,314 studied here (File S16) (Table 1).

We ran ExInAtor using protein-coding gene annotations, without stratification. Altogether, a total of 24 protein-coding drivers were identified at a false discovery rate cutoff of Q<0.1. Of these, 38% are CGC genes (indicated in red, Fig. 4A). This represents enrichment of 14-fold over random expectation (P<=0.0001, Chi-square with Yates' correction test). The most significantly enriched gene in this analysis is *TP53,* the most frequently mutated across cancers and identified in previous exome sequencing projects (16). *TP53* exons display an obvious and consistent enrichment of somatic mutations in both datasets, clustered in exons 4 and 7-11 (Fig. 4D). This *TP53* signal is observed in both Pancancer and multiple individual cancer types.

**Fig 4:**
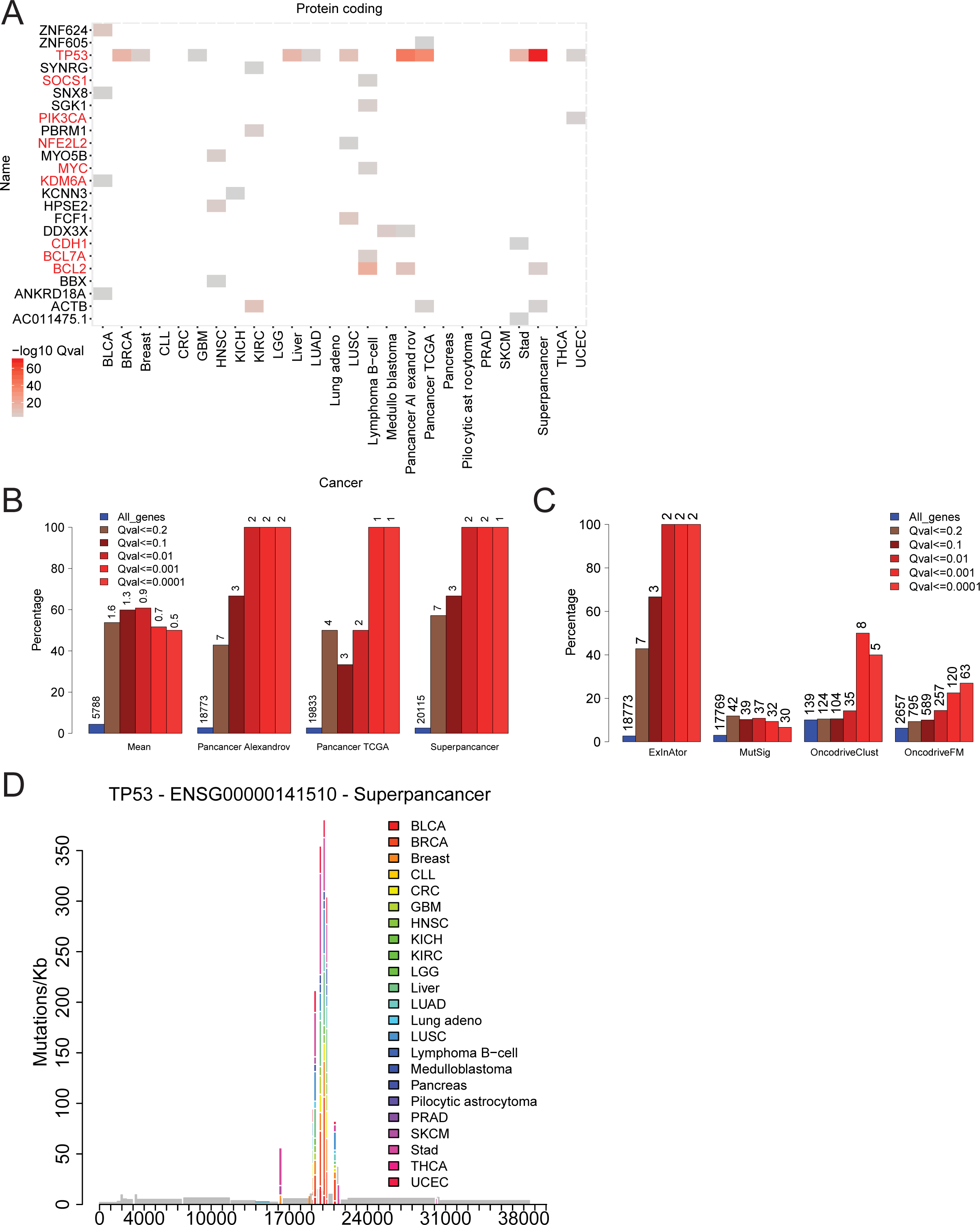
ExInAtor discovers known protein-coding drivers at high precision. (A) The −log10 Q-values of all candidates at Q≤0.1 cutoff in the Alexandrov and TCGA datasets. Gene names in red indicate known drivers belonging to the Cancer Gene Census (CGC). (B) The precision of ExInAtor predictions was estimated as the percent of predicted driver genes that also belong to CGC. Bars are coloured by the Q-value cutoff used, and the fraction of all known genes belonging to CGC is shown in blue as a reference. “Mean” displays the average overlap across all individual cancer types. The numbers above each bar indicate the total number of predicted driver genes at that cutoff. For example, in “Superpancer”, a total of three candidates are identified at a cutoff of 0.1, of which two (66%) belong to CGC. (C) Comparison of the performance of ExInAtor to other methods for protein-coding driver gene discovery, using the Alexandrov Pancancer dataset. Plot description as for Panel B. (D) The mutational profile of the TP53 tumour suppressor gene across all cancers. *x* axis indicates the position within the gene, *y* axis shows the mutation frequency.

Several of the 15 non-CGC genes identified have evidence for cancer roles: *ANKRD18A* in lung cancer (30), *DDX3X* and *PBRM1* in various cancers (31), *HPSE2* in thyroid carcinoma (32), *MYO5B* in gastric cancer (33). These findings suggest that ExInAtor precision may be better than implied by the analysis of CGC genes alone.

We examined the performance of ExInAtor, in terms of the percent of predicted genes that belong to CGC, at a series of Q value thresholds (Fig. 4B) (File S17). Shown are separate analyses for all cancer types (expressed as mean prediction per cancer), and various pancancer combinations. These show that, although the number of predicted genes are low, they tend to have far higher rate than that 2.7% expected by random chance, even at a Q value threshold of 0.1.

In summary, ExInAtor performs well in identifying known cancer related genes at high precision from a protein-coding training set ~10 times larger than CRL lncRNAs.

### ExInAtor is competitive with tools designed for protein-coding genes

Next we compared ExInAtor to a series of well-known pipelines for identification of protein-coding drivers: MutSig (17), OncodriveFM (22) and OncodriveClust (34). In side-by-side analyses on identical Alexandrov Pancancer data, we found that ExInAtor has low sensitivity (ie makes few predictions), but has excellent precision. In fact, its predictions contain a higher percentage of CGC genes than the other methods (Fig. 4C and File S18). For example, at a cutoff of Q<0.1, ExInAtor predicts 3 genes (of which 2 are known drivers), compared to 4 known drivers out of 39 for MutSig, 11 known drivers out of 104 for OncoDriveClust and 59 known drivers out of 589 for OncodriveFM (Fig. 4C). Furthermore, comparing the top 30 candidates detected at several cutoffs (File S19), the majority of genes detected by ExInAtor are also detected by at least one other method.

We also compared the four programs’ performance on real and simulated Pancancer data, displayed as Q-Q plots in Files S8 and S20. Again, ExInAtor performs relatively well: its predictions on true data mirror the expected distribution quite well, and true P values are smaller than for simulated data. ExInAtor predictions appear to be conservative, having a tendency for moderately deflated P values. In contrast, other methods tend to perform worse, being either strongly deflated (MutSig), inflated (OncodriveFM) or predicting less in true than randomised data (OncodriveClust). In summary, despite not employing any information from protein-coding sequence to inform its predictions, ExInAtor is surprisingly competitive with existing methods in the identification of coding driver genes. In particular, its predictions have low sensitivity (possibly many false negatives) but high precision (a high fraction of true positives). This lends weight to the accuracy of ExInAtor’s lncRNA predictions.

### LncRNAs are predicted as drivers at higher rates compared to coding genes

We were interested in the overall rates of prediction of lncRNAs and protein-coding genes, as well as their apparent tumour-specificity. Known driver genes are highly variable with respect to their tumour-type specificity. *TP53* mutations are found across a wide range of cancers, while other drivers are only mutated in single tumour types (16, 31). In this analysis, we detected no lncRNAs in more than one tumour (File S21). In contrast, two coding genes were discovered in two independent cancer types, while *TP53* was identified in no less than 9. Interestingly, a higher fraction of lncRNAs was predicted as driver genes than protein coding: 0.25% and 0.11%, respectively. These figures are likely to be strongly influenced by both the low sensitivity of ExInAtor discussed above and by the sparse data. In future, many more genes are likely to be identified in multiple cancers when deeper data is available. Nevertheless these findings suggest that lncRNA are mutated in cancer at a rate similar to, or higher than protein-coding genes.

### Novel and known driver lncRNAs share distinctive features of functionality

Returning to the driver lncRNAs identified by ExInAtor, we next asked whether any features distinguish these from other lncRNAs. Previous studies of lncRNA have used features such as evolutionary conservation and expression as proxies for functionality (35, 36). Furthermore, previous research on protein-coding cancer genes showed that their genes and their processed transcripts tend to be longer than average (37).

We compiled a series of features and, for each one, asked to what extent it differs between the CRL genes and all other lncRNAs. The full set of results, plotted by magnitude of difference and statistical significance, are shown in Figure 5A. It is clear that CRL genes are distinguished by a diverse range of features. They are transcribed from longer genes, and have longer mature transcripts (“exonic length”). They are under stronger evolutionary constraint: their promoters and exons are more evolutionarily conserved across a range of evolutionary distances. Their steady state RNA levels are higher and more variable across human tissues. Finally, they are also more likely to be proximal to a binding site of the P53 tumour suppressor. In contrast, there is no difference in genic or exonic GC content between CRLs and other genes.

**Fig 5:**
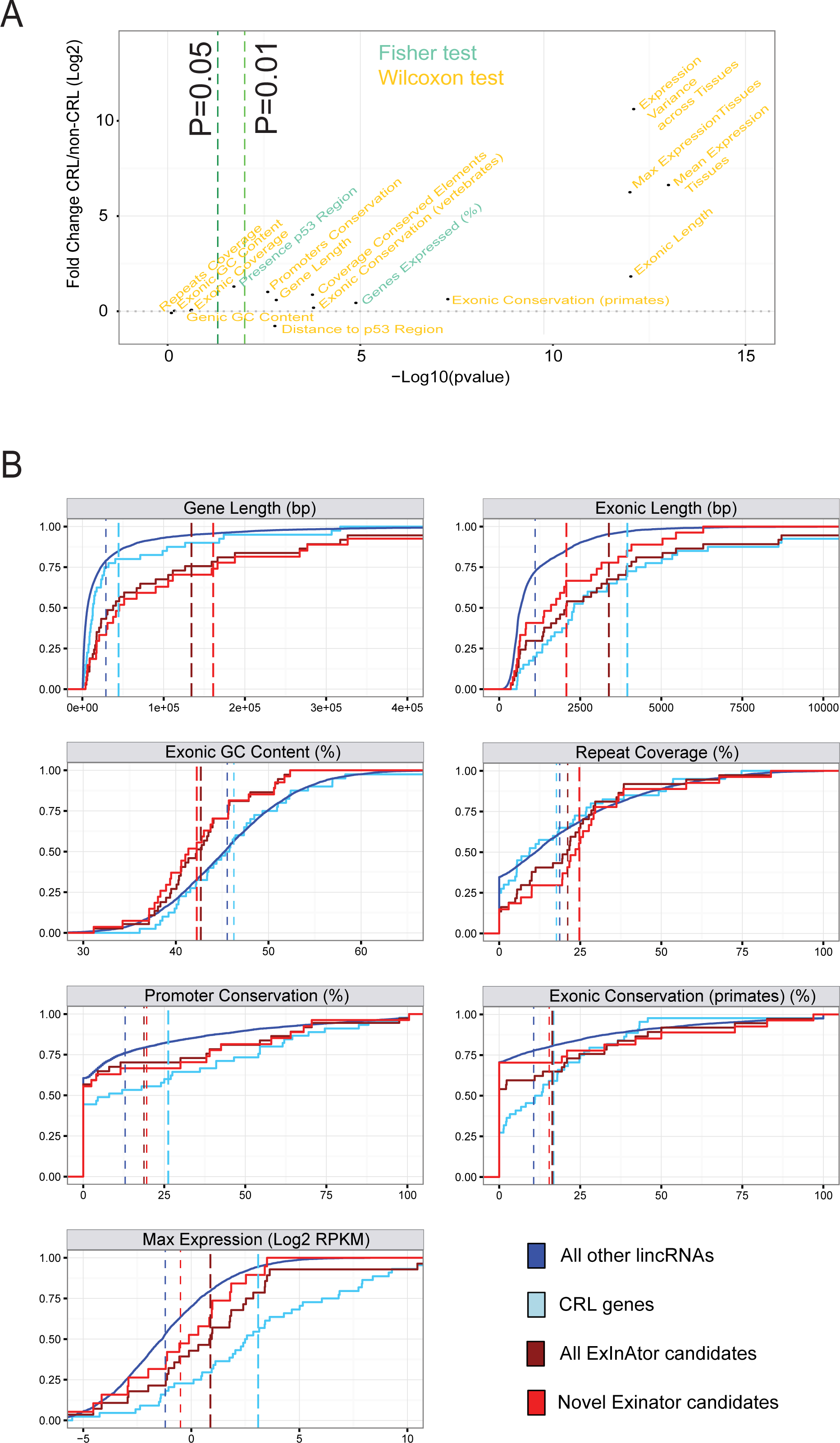
Features distinguishing cancer lncRNAs are also found in ExInAtor candidates. A) Identification of cancer lncRNA features by comparing literature-curated Cancer Related LncRNA (CRL) genes to others. Dots represent 16 features that were compared in CRL and non-CRL genes. *y* axis shows the log2 fold difference of CRL vs non-CRL means for the values of the given feature. *x* axis represents the P value obtained from the statistical test applied when comparing CRL and non-CRL. Features are coloured depending on whether they are discrete features, analysed by Fisher Exact test, or continuous features analysed by Wilcoxon test. B) Cancer lncRNA features in ExInAtor-predicted driver genes. Shown are cumulative distributions for seven selected features. Dashed vertical lines indicate the mean value of each group. Genes are grouped by: literature-reported cancer CRL lncRNAs, all ExInAtor candidates (both CRL and not)(“All candidates”), only novel ExInAtor candidates that are not included in CRL (“novel candidates”), and non-CRL non-candidates, being all other GENCODE lincRNAs. Candidates here were defined at a Q≤0.2. Groups significantly different from the latter at a threshold of P=0.05 (Wilcoxon test) are represented by a thick line.

Having established a series of cancer lncRNA-specific features, we asked whether these features are also present in ExInAtor candidate genes. We were particularly interested in whether novel candidates (ie non-CRL) share these characteristics, since this would represent an independent test for the value of ExInAtor predictions. Therefore we compared the features of three gene groups: CRL lncRNAs, all ExInAtor candidate genes, and novel ExInAtor candidates alone. These groups were compared to the null set of genes, represented by the entire set of remaining Gencode lincRNAs (“All other genes”).

In Figure 5B are shown the results across seven selected features. ExInAtor candidates, in common with CRL, have longer genes and transcripts than lincRNAs in general (P=4E-8, P=6E-4, respectively, Wilcoxon test). Surprisingly, and in contrast to CRL genes, ExInAtor candidates have significantly lower GC content (P=7E-3), and higher repetitive sequence content (P=0.03). Finally, for features of evolutionary conservation of both promoter and exon, in addition to steady-state RNA levels, we find that novel candidates display a similar trend as CRL genes, although these do not reach statistical significance (P>0.05). In summary, and pending future replication with larger gene sets, it appears that novel ExInAtor predicted cancer genes share a number of distinguishing features with known cancer lncRNAs, consistent with being *bona fide* driver genes.

## Materials and Methods

### Gene annotation and filtering

The GENCODE v19 lncRNA catalogue was downloaded in GTF format from (www.gencodegenes.org) (4, 38), and comprises 13,870 genes. A number of filtering steps were applied to this list. First, only intergenic genes (having no transcripts overlapping protein-coding genes on the opposite strand, or within 10 kb at their closest point on the same strand) were retained (6, 308). Second, any lncRNA gene with transcripts of uncertain protein-coding potential were removed, leaving 5,887 genes (see below for details). Third, we included several cancer-related lncRNAs from the scientific literature, resulting in a final set of 5,914 lncRNA genes (Table 1 and Files S3 and S4). Note that literature genes may violate the two filters above, but must have a GENCODE identifier. This set of filtered lncRNAs is used throughout.

The protein-coding gene catalogue was also obtained in GTF format from GENCODE v19 (38). From this annotation, all genes with biotype “protein-coding” were selected, resulting in 20,314 genes and 145,518 transcripts. Finally, all transcripts not having biotype “protein-coding” were removed, reducing the transcripts to 81,702. (Table 1 and File S16).

### Somatic mutation data curation

Whole-genome cancer somatic mutations were obtained in BED format from two sources: 10 cancers described in Alexandrov et al (20), and 14 cancers from TCGA (1). In addition, we included an additional dataset of 100 stomach adenocarcinoma (STAD) with the Alexandrov dataset (39), resulting in an original set of 22,877,059 mutations. Only single nucleotide somatic mutations and indels of length 1 were retained (97.7% of the total somatic variants), hereafter referred to as “mutations”. AML and ALL cancers from the Alexandrov dataset were removed due to their low number of genomes and mutations. Statistics on the remaining cancers can be found in Table 2. Both mutation datasets were prefiltered in order to remove possible misannotated germline SNPs. First, any mutations identical to an entry in dbSNP 146 “common” (>1% frequency) were removed, leaving 22,128,594 mutations (96.7%). Second, any recurrent mutations, having the same nucleotide change observed in the same location more than once, were collapsed and treated as a single event, resulting in a final set of 20,837,263 mutations (91.1%).

### Assessing the protein-coding potential of lncRNA

All GENCODE v19 lncRNA transcripts were tested for protein-coding potential with CPAT (40) at default settings. Any gene having one or more transcripts predicted to be protein-coding (coding potential >= 0.364) was removed from further analysis.

### ExInAtor design

ExInAtor requires eight mandatory inputs: (1) a gene annotation in GTF format containing information on genes and exons to analyse (transcript information is ignored), (2) a catalogue of mutations in BED format, (3) the number of individual genomes or samples represented by the BED file, (4) the output folder destination, (5) a file with two columns showing the name of each chromosome and its nucleotide length, (6) a gene annotation in GTF format containing information on genes and exons of the whole genome (transcript information is ignored), (7) FASTA file of the whole genome and (8) a file containing all the possible trinucleotides. Optional inputs are: (1) a minimum number of exonic and/or (2) background mutations that each gene must have to be analysed, (3) the number of CPU cores to use in the analysis and (4) the extension length of the background region that includes all introns

The ExInAtor workflow can be divided into the following steps: exon and background definition, mutations mapping, sub-sampling of background region, gene filtering by mutation counts and statistical analysis (Fig. 1).

Exon / Background definition (File S1-A-B-C): The full set of exons from all transcripts belonging to a gene are merged. The remaining genic space is then defined as background, which is extended to both sides of the gene according to the window length parameter. In the present study, this value was set at 10 kb throughout. Regions overlapping exons from any other gene are removed from this background region. The coordinates of non-overlapping exons and background regions are saved in BED format. The total exonic and background nucleotide length is calculated.

Mutations mapping (File S1-D): Mutations are mapped to exons and background regions, then counted. Despite in this study we collapsed the recurrent mutations in only one, if two or more mutations fall in the same position they are counted separately.

Sub-sampling of background region (File S1-E-F): The trinucleotide content of the exonic and background regions are calculated. Then, regions of identical size and trinucleotide composition to the exonic region are sampled from the background region. This is performed sequentially, without replacement, until it is impossible to continue. At every iteration, the sampled positions are added to a new background region, along with their associated mutations. In this way, a new background region of maximal size and identical composition to the exonic region is assembled for every gene.

Gene filtering by mutation counts: Mutation data are sporadic and of low density, potentially resulting in inflated P values. To avoid this, ExInAtor accepts a user-defined minimum number of exonic and background mutations, below which lncRNAs will not be considered. These cutoffs may be defined by the user, with the default filter (used in the present study) discarding genes with less than 1 exonic mutations or 1 background mutations.

Statistical analysis: Statistical enrichment of exonic mutations is determined using the hypergeometric test (Fig. 1B). The following contingency table is compiled for each gene, with the total exonic and background lengths, N and n respectively:

M = number of exonic positions mutated
N − M = number of exonic positions not mutated
m = number of background positions mutated
n − m = number of background positions not mutated

This is the starting point for calculations of statistical significance of enrichment of exonic mutations using the hypergeometric distribution, which describes the probability of obtaining a given number of successes in a given number of draws without replacement from a finite population of a specific size. It is important to note that the positions corresponding to each genome are counted independently, meaning that the total gene length *N* is defined as gene length multiplied by the number of genomes. *n* is treated similarly. Statistical significance is estimated for a gene to have that many or more exonic mutations, then are corrected for multiple hypothesis testing using the Benjamini-Hochberg procedure, which controls the False Discovery Rate (FDR), here indicated by “Q”.

ExInAtor returns the input gene list with mutation counts and associated exonic enrichment Q-values. The latest ExInAtor version is freely available for download here:https://github.com/alanzos/ExInAtor/

### Creation of a simulated mutation dataset

Two distinct methods were used to create trinucleotide-aware simulations of tumour mutations. In the first method (“Fixed window reassignment”), the genome was divided into fixed partitions of 50 kb. Mutations were randomly assigned to another genomic location with the same reference trinucleotide and surrounding nucleotides for substitutions and indels, respectively. In the second method (“Sliding window reassignment”), a 50 kb window is centred on each individual mutation. The mutation is then reassigned to another position with identical reference trinucleotide within its window. These simulations, while maintaining approximately the same number of single nucleotide substitutions and indels of the original Alexandrov dataset as well as the same mutation trinucleotide signature, constitute neutral datasets that are not expected to be enriched in cancer related lncRNA.

### Visual inspection and validation of candidates’ mutations

To verify the quality of the mutation calling, we visually validated 12 single somatic mutations from 4 candidates. First, we downloaded a SAM file of the surrounding regions of each mutation (+/− 2kb) with the BAM Slicer tool from CGHUB (https://cghub.ucsc.edu/); then we opened those files with IGV to check the reads supporting the mutations (File S22) (41).

### Comparison of cancer features

We obtained the list of lncRNA genes proximal to cancer-related germline SNPs from Table S5 of (29). The enrichment of indicated lncRNA genesets with respect to these genes were assessed by contingency table analysis using Fisher’s Exact text. For the analysis of CNVs, the set of regions were obtained from Table S3 of the same paper, and statistical enrichments were calculated similarly. Only data from cancers corresponding to those in the present study were considered.

### Comparison of lncRNA features

To assess which features may distinguish cancer lncRNAs, we collected different genomic and expression data for all the genes and divided them into the four groups of interest:

1. non-CRL non-candidates (non-CRL gene list excluding ExInAtor discoveries at a Q≤0.2, “All other lincRNAs”)
2. CRL genes
3. all ExInAtor candidates (discovered at a Q<0.2)
4. novel ExInAtor candidates (candidate genes that do not appear in the CRL list).

For each feature we compared all groups to non-CRL non-candidates. Statistical tests were performed using R. Features were compiled from the following sources:

#### Gene Sequence

Gene sequence features were calculated based on Gencode v19 annotations. Exonic regions of each gene were defined as the projection of all exons from the union of its transcripts. Promoter regions of each gene were defined as a window of +/- 100 nucleotides from the reference transcription start site (TSS).

#### Conservation

PhastCons scores from vertebrate and primate species alignments and PhastCons Elements from vertebrate, mammals and primate species alignments were downloaded from UCSC Genome Browser. Two separate analyses were performed, using either base-level scores, or conserved element regions. We separately computed the average exonic base-level conservation score of each gene for primates and vertebrates PhastCons scores. We merged conserved elements annotations from primate, mammal and vertebrate species alignments and intersected these regions with promoters and exonic regions. We then computed the percent of nucleotides (from promoters or exonic regions) covered by conserved elements for each gene.

#### Repeat Elements

We downloaded the 2013 version of RepeatMasker human genomic repetitive element annotations and converted it to BED format. These annotations were intersected with exonic regions of lncRNAs. For each gene we calculated the percent of exonic nucleotides overlapping repetitive elements.

#### Tissue Expression Analysis

We extracted tissue expression values for 16 human tissues from Human Body Map (HBM) RNAseq data, downloaded from ArrayExpress under accession number E-MTAB-513. These data were used to quantify Gencode v19 genes using the GRAPE pipeline (42). Considering only genes that are expressed (RPKM>0) at least in one tissue we described the mean, the maximum and the variance of RPKM expression values across tissues. The percent of expressed genes for a given group represents the total number of genes that are expressed at least in one tissue compared to the total number of genes of the given group.

#### P53 analysis

We obtained ChIP data for p53 binding sites from (28). Binding maps from the two available timepoints were merged. We attempted to assess a possible link between cancer driver lncRNAs and p53 binding site regions in two different ways. We first analysed whether the position of CRL genes in the genome tend to be closer to p53 binding site regions compared to non-CRL genes. To this aim, we calculated the nucleotide distance from the promoter of the gene (defined as explained before) to the closest p53 binding site region for all CRL and non-CRL genes. As an alternative, we compared the probability of finding a p53 binding site close to a TSS for CRL and non-CRL genes: for each we counted how many genes out of the total contain at least one predicted p53 binding site region within a window of 100kb, centred on the TSS.

## Discussion

Here we have presented ExInAtor, to our knowledge the first method specifically designed to identify cancer driver lncRNAs from tumour genome cohorts. ExInAtor aims to address the unique opportunity of comprehensively discovering cancer driver lncRNAs within and across tumour types using mutation data generated by projects such as TCGA and ICGC.

We have presented the results of scans across the two most substantial tumour genome sequencing cohorts presently available, the Alexandrov and TCGA datasets, altogether comprising more than 1000 genomes from 23 cancer types. In addition to successfully retrieving at nine known protein coding drivers (38% of total predictions) and six published cancer-related lncRNAs (40% of predictions), we identify for the first time a total of nine novel lncRNA driver genes at low false positive rates (0.1 FDR). These novel candidates share with known cancer lncRNAs a series of features including evolutionary conservation, normal tissue expression and gene length. They also tend to be proximal to germline cancer SNPs and have increased probability of lying in CNV regions, lending weight to their association with tumourigenesis. Together these observations lend weight to the idea that ExInAtor predicts *bona fide* driver lncRNAs. The true test of these predictions must await experimental validation in cell lines and animal models.

The distinguishing features of cancer-related lncRNAs are reminiscent of similar findings for protein coding genes (37). Evolutionary conservation and high steady-state RNA levels are generally interpreted in this context as evidence for functionality of lncRNAs (35, 36). The significance of other features is less clear, and we should be careful to consider possible non-biological factors. In the case of gene length, it is likely that ExInAtor has greater statistical power for longer genes, possibly explaining the significantly elevated lengths of known and novel candidates. Furthermore, it is likely that the annotated length of lncRNAs is correlated with their expression, since higher expressed genes have more supporting ESTs and cDNAs, and hence are more complete.

Other observations were unexpected: the exons of novel candidate drivers have elevated repetitive content and reduced GC content. Furthermore, and in contrast to the above, these features are not shared with known CRL driver genes. It is unclear whether this reflects technical artefacts of the analysis, or a genuine biological insight. We can think of no bias in ExInAtor, or the cancer mutation datasets, that may favour gene models with these properties, although it is entirely feasible. On the other hand, transposable elements have been linked to both cancer (43, 44) and lncRNA functionality (45). It is attractive to hypothesise that repeat-rich lncRNAs play roles in tumourigenesis and are preferentially mutated during this process. Further study will be required to establish the significance of these findings.

At present, our understanding of how lncRNA function is encoded in sequence motifs and structures is limited (19). Consequently, advanced approaches for scoring the functional effect of mutations, such as those used for protein sequences, are unavailable. Nevertheless, future improvements to ExInAtor may include information on RNA structures, protein binding sites, post-transcriptional processing and evolutionary conservation to weight mutations based on their likely impact on lncRNA function. Furthermore, more sensitive statistical methods employing information on mutation clustering and cancer-specific mutational signatures will likely improve predictions.

We expect that future studies will yield many more candidate lncRNAs than produced here. Although the datasets used represent a large proportion of all presently available tumour genomes, future projects will likely be larger and produce mutation calls of better quality. For example, the upcoming PCAWG project will likely produce several fold more genomes than used here, and with more sophisticated mutation calling (1, 46).

The increasing scale of cancer genome projects will place a growing emphasis on computational efficiency. One of the benefits of ExInAtor is its ability to handle data with complex trinucleotide biases uses a simple subsampling algorithm, and without any functional impact predictions. This simplicity has the unintended benefit that ExInAtor is capable of identifying protein-coding drivers with precision comparable to the best methods. Another outcome is that ExInAtor makes very low computational demands: analyses for this paper were executed on a workstation running Intel Core i7 processors. 25 minutes were required to analyse protein coding genes in Superpancancer (the largest dataset tested here) using a single core and 2,050 MB of RAM. It required just three minutes to analyse Pilocytic astrocytoma with six cores and 648 MB of RAM. Together, these features make ExInAtor suited to future, large-scale cancer genome sequencing projects.

## Acknowledgements

The authors wish to thank Núria López-Bigas for her support throughout the project. We also thank Marta Melé (Harvard University) for insightful discussions, and Maite Huarte (CIMA) for insightful discussions and providing P53 binding maps, and Darek Kedra (CNAG) for the help with the visual inspection of the somatic mutations. We especially thank Erik Larsson (University of Gothenburg) for sharing TCGA mutation calls ahead of publication. We thank Romina Garrido (CRG) for administrative support. We acknowledge support of the Spanish Ministry of Economy and Competitiveness, ‘Centro de Excelencia Severo Ochoa 2013-2017’, SEV-2012-0208. R.J. is supported by Ramón y Cajal RYC-2011-08851 and Plan Nacional BIO2011-27220. A.L. is supported by pre-doctoral fellowship FPU14/03371.

## Author contributions

R.J. conceived the project, and supervised with advice and suggestions of R.G. Primary development of the tool was carried out by A.L., as well as the creation of one of the simulated datasets. Statistical and technical assistance for running analysis were provided by F.R and E.P., respectively. A.L. and J.C. performed the secondary analysis. L.M. created one of the simulated datasets, executed the analysis of MutSig, OncodriveFM and OncodriveClust on protein coding genes. R.J., A.L. and J.C. drafter the manuscript and prepared the figures and supplementary material. All authors read and approved the final draft.

